# OncoScape: Exploring the cancer aberration landscape by genomic data fusion

**DOI:** 10.1101/041962

**Authors:** Andreas Schlicker, Magali Michaut, Rubayte Rahman, Lodewyk FA Wessels

## Abstract

Although large-scale efforts for molecular profiling of cancer samples provide multiple data types for many samples, most approaches for finding candidate cancer genes rely on somatic mutations and DNA copy number only. We present a new method, OncoScape, which, for the first time, exploits five complementary data types across 11 cancer types to identify new candidate cancer genes. We find many rarely mutated genes that are strongly affected by other aberrations. We retrieve the majority of known cancer genes but also new candidates such as *STK31* and *MSRA* with very high confidence. Several genes show a dual oncogene-and tumor suppressor-like behavior depending on the tumor type. Most notably, the well-known tumor suppressor *RB1* shows strong oncogene-like signal in colon cancer. We applied OncoScape to cell lines representing ten cancer types, providing the most comprehensive comparison of aberrations in cell lines and tumor samples to date. This revealed that glioblastoma, breast and colon cancer show strong similarity between cell lines and tumors, while head and neck squamous cell carcinoma and bladder cancer, exhibit very little similarity between cell lines and tumors. To facilitate exploration of the cancer aberration landscape, we created a web portal enabling interactive analysis of OncoScape results.

## INTRODUCTION

Cancer is one of the biggest public health problems with an estimated number of 14.1 million new diagnoses and 8.2 million deaths worldwide in 2012 according to GLOBOCAN (http://globocan.iarc.fr/Pages/fact_sheets_cancer.aspx). Large-scale initiatives such as The Cancer Genome Atlas (TCGA) (1) and the International Cancer Genome Consortium (ICGC) (2) have been established specifically with the aim to determine the mechanisms underlying the development and progression of all major cancer types. To this end, large numbers of tumors and matched normal samples of different cancer types were extensively molecularly characterized. Using these data, candidate cancer genes were identified, most often relying on significantly elevated somatic mutation rates (3) rather than integration of different data types. Several published analyses combined somatic mutations across cancer types in order to increase the power for discovering potential cancer genes (4–6). However, even approaches relying on partially overlapping somatic mutation data sets (5, 6) show limited agreement in their results. A few other studies integrated mutation and DNA copy number data (7) or gene expression, copy number and DNA methylation data (8). Recently, Sanchez-Garcia and colleagues developed a new method that first identifies regions with recurrent copy number gain and then prioritizes the contained genes using data from exome sequencing, shRNA screens and gene expression (9).

Cancer cell lines are important tools for basic cancer research and efforts are currently underway to molecularly characterize large cell line panels representing a highly diverse set of cancer types (10–12). Since cell lines grow in an artificial environment, it is often unclear how well they represent primary tumors. Recent studies concluded that colorectal cell lines represented tumors well (13), while some ovarian lines showed features uncharacteristic of primary tumors (14) and bladder cancer cell lines also differed significantly from primary tumor samples [].

In this study, we describe OncoScape, a new method for integrating gene expression, somatic mutations, DNA copy-number and methylation as well as data from shRNA knock-down screens. To the best of our knowledge, this is the first approach to integrate such diverse data types for identifying cancer gene candidates. Applying OncoScape to 11 cancer types, we provide a more accurate picture of the landscape of molecular aberrations of human tumor samples. We specifically aim to identify novel candidate cancer genes that are aberrated in several cancer types and exhibit aberrations in different data types. Our method analyzes data for each cancer type separately and then aggregates the results on a gene level across cancer types. In contrast to most previous analyses, we classify candidate cancer genes as potential oncogenes or tumor suppressor genes based on the observed types of aberrations. Our results correlate well with the literature in the sense that known oncogenes and tumor suppressor genes are scored highly by OncoScape. Importantly, we identify many highly aberrated genes in different cancer types that have not been implicated in cancer so far. As it is known that aberrations cause (in)activation of specific pathways in cancer (15, 16), we analyzed aberrations across pathways, which shows a birds-eye view of the aberration landscape and facilitates the identification of functional themes in each cancer type. We performed the same analysis for cell lines of ten cancer types providing the most comprehensive comparison of the aberration landscapes of cancer cell lines and primary tumor samples to date. We identified several cancer types, such as ovarian, breast and colon cancer, for which primary tumors and cell lines show high similarity. However, in other cancer types, especially bladder cancer and head and neck squamous cell carcinoma, cell lines exhibited only limited similarity with tumor samples, emphasizing the importance of carefully selecting *in vitro* model systems.

## MATERTIALS AND METHODS

### Data acquisition and accessibility

Gene expression, DNA copy-number and methylation data for primary tumor samples and matched normal samples were obtained from data freeze 4.7 of the TCGA Pan-Cancer (1) project entry hosted in Sage Bionetworks’ Synapse (Synapse accession doi:10.7303/syn300013). In all analyses, only tissue-matched normal samples were used. If a dataset contained one primary tumor sample and a recurrent or metastasis sample of the same patient, only the primary sample was retained. If several primary samples were available for one patient, we removed all samples from the dataset. Somatic mutations for TCGA tumor samples were also downloaded from Synapse (Synapse accession doi: 10.7303/syn1710680.4). We divided mutations in the colorectal dataset into separate sets for colon and rectal cancer. We obtained shRNA knock-down data from Project Achilles (12) on 20 June 2013. Gene expression, mutation and DNA copy-number data for cancer cell lines were downloaded on 12 March 2012 from the CCLE web site. Cell lines were assigned to cancer types based on their tissue type annotation (Tables S8 and S9). In total, we matched 327 cell lines to one of ten cancer types included in our analysis; there were no rectal cancer cell lines in the dataset. For DNA copy number and gene expression data, we corrected for systematic differences between CCLE and TCGA utilizing the ComBat function in the sva package (version 3.6.0). For DNA copy number data, we applied ComBat on the combined matrix containing all CCLE and TCGA samples correcting for the batch variable (CCLE or TCGA) while taking into account tissue type and disease state (cancer or normal) as outcome of interest. Batch correction for gene expression was performed in the same way but separately for tissue types with RNA sequencing-or chip-based expression data. For the analysis of colorectal subtypes, we utilized the random forest subtype labels from (17). We did not use shRNA data for this analysis as no clear assignment of cell lines to subtypes was available. Pathway definitions were taken from the Kyoto Encyclopedia of Genes and Genomes (18) on 25 February 2014 using version 1.0.1 of the KEGGREST package. Gene IDs were translated using version 2.16.0 of the biomaRt package (19). All results of our analyzes are publicly accessible in Synapse (https://www.synapse.org/, Synapse accession doi:10.7303/syn2518467).

### OncoScape

Data processing and visualization was performed using the R statistical programming environment (20). We implemented a new package, named OncoScape, for data processing and prioritization of candidate cancer genes. Source code of the package and all other scripts used for data handling and visualization can be freely downloaded from Github at https://github.com/andreas-schlicker/OncoScape. The package provides functionality to assess alterations of single genes for each data type and each cancer type individually (Figure 1). Tumor samples are compared with normal samples to identify significant differences. If a gene is found to be altered, this gene receives a score of 1 for this data type and else a score of 0. Details on the scoring for each data type are given below. Activating and inactivating alterations are both scored independently for each gene, and the sums of the activating and inactivating aberrations yielded an oncogene score and a tumor suppressor gene score, respectively. Additionally, we calculated the difference between oncogene score and tumor suppressor gene score, referred to as overall score, and the sum between oncogene and tumor suppressor scores, referred to as aberration score. Genes were then ranked based on one of these scores to be classified as potential new oncogene or tumor suppressor gene. Pathway alteration scores were calculated by averaging scores for all genes assigned to the same pathway. Aberrations in cancer cell lines were assessed by comparing the cell lines with normal samples available from TCGA using the same approach as for tumor samples.

**Figure 1:**
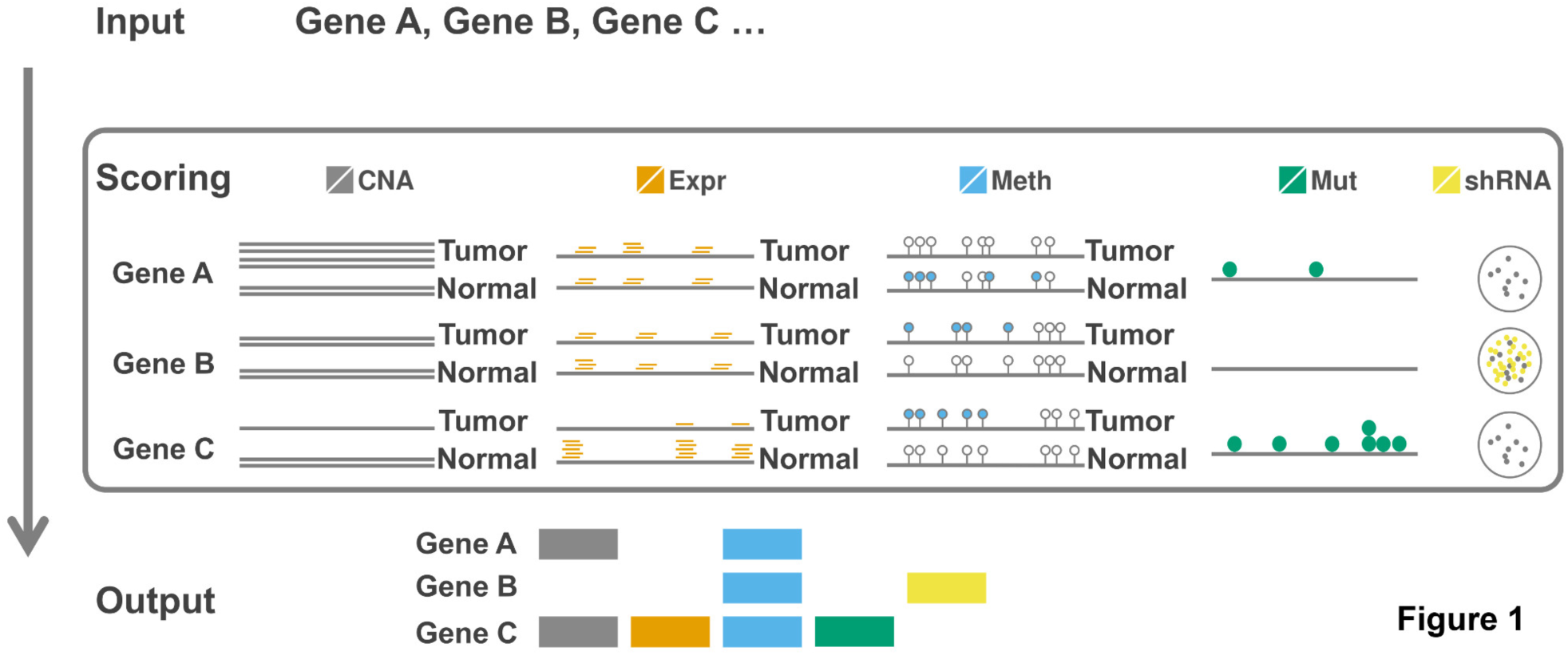
Schematic overview of the OncoScape prioritization method and the data types used. The input to the method consists of a list of genes scored for aberrations in five data types. Aberrations in copy number (CNA), gene expression (Expr) and DNA methylation (Meth) are identified as statistically significant differences between tumor and normal samples. Somatic mutations (Mut) are scored based on the type of mutations and their clustering along the gene (clustered mutations contribute to the oncogene score, while broadly distributed mutations contribute to the tumor suppressor score). Gene knockdown (shRNA) results are assessed using the change in cell line growth before and after knockdown. Filled boxes in the output table indicate aberrations identified for individual genes in each data type. Inactivating aberrations count for the tumor suppressor score while activating aberrations count for the oncogene score. This analysis is executed for each cancer type separately.

### Gene expression analysis

Normalized gene expression data for tumor and normal samples, either from an Illumina sequencing platform or Agilent arrays depending on availability for each cancer type, were obtained from TCGA. We compared expression levels for each gene between tumors and matched normal samples using paired Wilcoxon tests and corrected nominal p-values using the Benjamini-Hochberg procedure. If a gene was significantly (FDR < 0.05) differentially expressed (lower or higher) in tumor samples, it received a +1 towards tumor suppressor or oncogene score, respectively.

### Copy-number analysis

Segmented DNA copy-number data for tumor and normal samples were obtained using Affymetrix SNP6 arrays by TCGA. We compared log-ratio copy number values between tumor samples and matched normal samples using paired Wilcoxon tests and corrected p-values using Benjamini-Hochberg’s procedure. We required genes to have a mean copy number difference larger than 0.1 between tumors and normal to prevent small effects becoming significant because of a large number of samples. For genes with significant difference in copy-number (FDR < 0.05), we calculated the Spearman correlation between copy-number and gene expression. A gene was scored as potential tumor suppressor or oncogene if 1) its copy-number value in tumor samples was significantly lower or higher than in normal samples and 2) the copy-number was significantly (FDR < 0.05) positively correlated with gene expression across the tumor samples.

### DNA methylation analysis

We analyzed DNA methylation data in a probe-wise fashion. For each probe, we compared methylation values in tumor and tissue-matched normal samples using unpaired Wilcoxon tests and corrected p-values using the Benjamini-Hochberg procedure. For each probe with a mean difference of methylation beta valus greater than 0.1 between tumors and normals and the difference being significant (FDR < 0.05), we computed the Spearman correlation between methylation level and expression of the genes annotated to that probe according to the Illumina annotation across all tumor samples. Correlations with FDR < 0.05 were regarded as significant. A gene was scored as tumor suppressor gene if 1) at least one associated methylation probe in the gene body exhibited significantly lower methylation in tumors and 2) the methylation was positively correlated with gene expression, or if at least one other associated methylation probe showed gain of methylation in tumors and gene expression was negatively correlated. In contrast, a gene was scored as oncogene if 1) at least one probe in the gene body showed gain of methylation and 2) was positively correlated with expression or any other associated probe exhibited lower methylation in tumors and methylation was negatively correlated with gene expression.

### Mutation analysis

Mutations were scored according to the 20/20 rule published by Vogelstein and colleagues (21), including classification of mutations into oncogene or tumor suppressor mutations, respectively, and the applied cutoffs. We divided mutations according to their classification into oncogene mutations (missense mutations and in frame deletion/insertions) and tumor suppressor mutations (frame-shift deletions/insertions, nonsense mutations and splice site mutations). Then, we calculated an oncogene mutation rate (OGMR) and a tumor suppressor mutation rate (TSMR) for each gene. The OGMR was defined as one minus the number of distinct oncogene mutations divided by the total number of mutations. The TSMR was defined as distinct tumor suppressor mutations divided by the number of total mutations. As suggested in [], genes with an OGMR > 0.2 and TSMR < 0.05 were scored as oncogene. A gene was scored as tumor suppressor if its TSMR > 0.2 or if the OGMR > 0.2 and the TSMR > 0.05. We required a minimum of five oncogene or tumor suppressor mutations in order to score a gene as oncogene or tumor suppressor, respectively.

### Analysis of shRNA knock-down screens

Project Achilles assessed cell viability after knocking down genes using different shRNA hairpins. In order to minimize off-target effects, Project Achilles integrated knock-down results of several hairpins targeting the same gene into so called “gene solutions”, providing a cell viability score for each gene and cell line combination. We used only genes for which only a single gene solution was provided. For each gene, we derived a distribution of viability values across all cell lines. A gene was scored as potential oncogene for one cancer type if at least 25% of the cell lines for this cancer type had a knock-down viability score that was lower than the 25th percentile across all cell lines. If at least 25% of the cell lines had a knock-down viability score greater than the 75th percentile for that gene, it was scored as potential tumor suppressor gene.

### Estimation of affected samples

In order to estimate which samples were affected by alterations in gene expression, DNA copy number or methylation of a specific gene, we first obtained the distribution of differences between tumor and normal samples. For paired analyses, we simply subtracted the value of the matched normal samples from the value of the tumor samples. For unpaired analyses, we subtracted the average value across all normal samples from the values of the tumor samples. A sample was called affected if its data value was more than one standard deviation away from the mean value of the distribution of differences.

## RESULTS

### Prioritization method

In order to obtain a comprehensive characterization of the molecular aberration landscape of different cancer types, we developed OncoScape, an algorithm integrating gene expression, DNA copy number, DNA methylation and somatic mutation data, as well as shRNA knock-down screens. Figure 1 provides a schematic overview of the workflow - a detailed description is given in the Methods section. Briefly, OncoScape prioritizes genes as potential oncogenes or tumor suppressor genes by identifying molecular aberrations based on a comparison of tumor and normal (same tissue) samples. For each cancer type, we identify genes whose mRNA expression, DNA copy number or methylation patterns differ significantly between tumor and tissue-matched normal samples. Patterns of somatic mutations along the gene and results of shRNA knock-down screens provide additional evidence. Calling aberrations for each gene and cancer type separately, enables us to identify cancer type-specific aberrations. Aberrations in the different data types are weighted equally; activating aberrations contribute towards an oncogene score and inactivating aberrations towards a tumor suppressor score. These scores are used to rank genes per cancer type, with higher scores signaling higher confidence in the prediction. Importantly, we primarily focus on finding genes with aberrations in different data types rather than those with only one type of aberration. Furthermore, we calculate a combined score as the difference between the oncogene and tumor suppressor scores.

### The global aberration landscape of tumors

The main goal of our study was to provide a comprehensive map of the global aberration landscape across 11 human cancer types. To this end, we prioritized all genes in the human genome taking into account five different data types (Table 1). Figure 2a summarizes the number of genes with aberrations in the different cancer and data types. Across cancer types, the number of affected genes was similar for activating (oncogene-like) and inactivating (tumor suppressor-like) aberrations but highly dependent on the data type. Consistent with previous analyses, relatively few genes were frequently mutated in each cancer type (5), while aberrations in DNA methylation were found at a much higher frequency (8). Interestingly, only glioblastoma (GBM), colon adenocarcinoma (COAD) and kidney renal clear cell carcinoma (KIRC) had appreciable numbers of genes with statistically significant expression changes. Considering only one data type, most genes were found to be aberrated in at least one cancer type (Table S1). However, only 1196 genes had either activating or inactivating aberrations in at least three data types in at least one cancer type. Few genes were aberrated in four data types in some cancer types (Table S2), including the well-known tumor suppressor genes *RB1, BRCA2* and *PTEN.*

**Figure 2:**
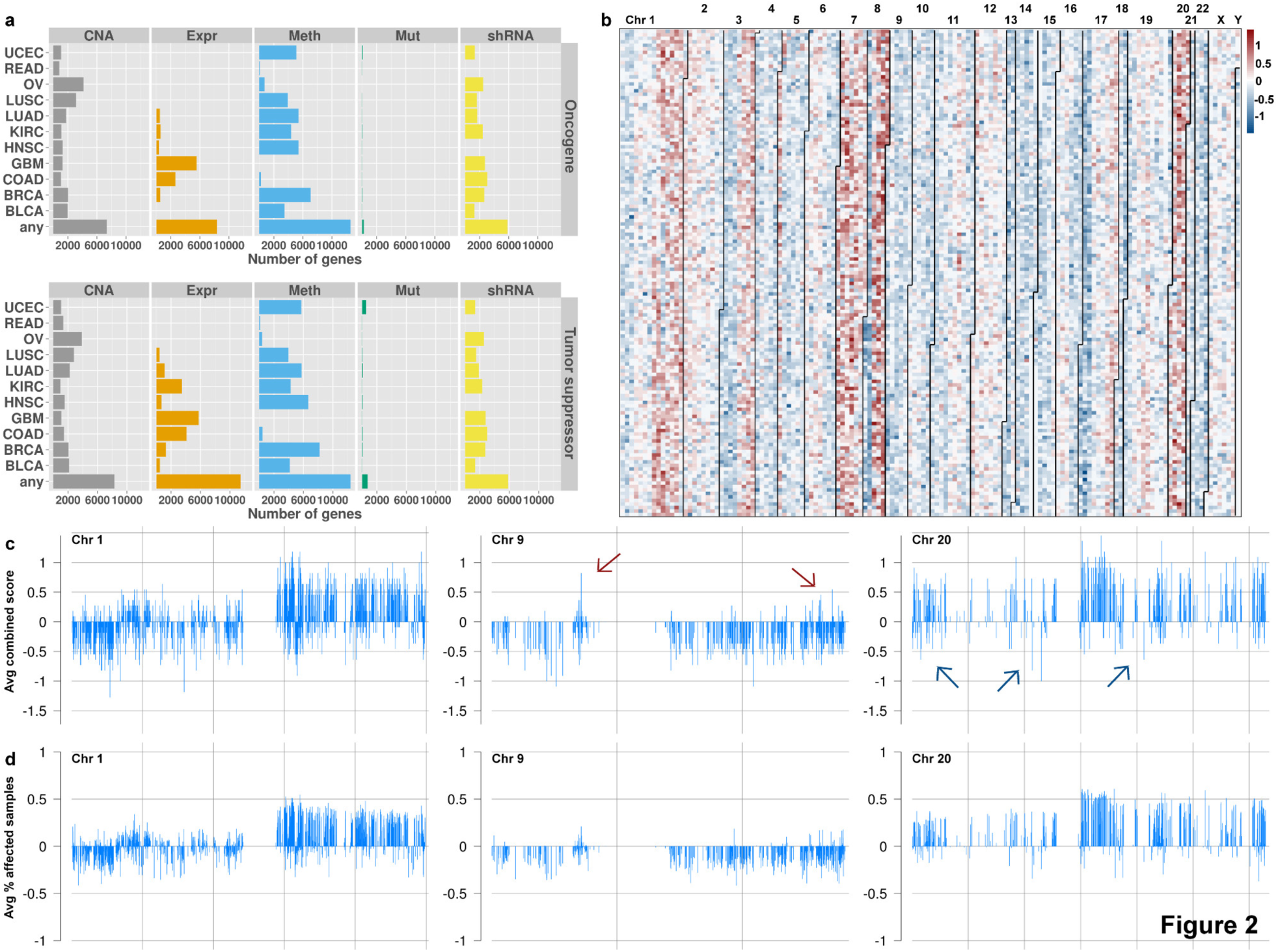
Overview of the molecular aberration landscape. (a) Number of genes with oncogene-like and tumor suppressor-like aberrations in the different data types across all cancer types. Data types are depicted in panels and cancer types in rows. The row labeled “any” indicates the number of genes with an aberration in any cancer type. (b) Two-dimensional overview of prioritization scores averaged across all 11 cancer types. Genes are plotted column-wise sorted according to chromosomal location, starting with the telomere of Chromosome 1 p in the upper left corner. Lines delineate the different chromosomes and numbers indicate chromosome names. Potential oncogenes have positive scores and potential tumor suppressor genes have negative scores. (c, d) Depiction of the average combined scores and average percentage of affected samples across Chromosomes 1, 9 and 20. Red and blue arrows point to predicted oncogenes and tumor suppressor genes, respectively, which are targeted by focal events. Panel (c) depicts the average prioritization scores for the genes; panel (d) shows the average percentage of samples affected by aberration in these genes. For clarity, percentages of samples affected by inactivating alterations are depicted as negative numbers.

Arranging genes by chromosomal location revealed large aberration patterns corresponding to chromosome arms or whole chromosomes (Figure 2b). Genes located on Chromosome 1p, for instance, exhibited primarily inactivating aberrations across different cancer types, while genes on Chromosome 1q were mostly targeted by activating aberrations (Figure 2c). Investigating contributions of individual data types showed that this pattern was primarily caused by large-scale DNA copy number changes (Figures S1-S5). However, in these large regions, specific genes had higher scores than the surrounding genes showing that these genes are hit by different types of aberrations. This suggests that these genes could be the targets of such large-scale copy number events while the remaining genes in these regions have a passenger role (Figure 2c,d). Chromosome 9, for example, largely targeted by inactivating alterations, contained individual genes showing activating aberrations. Across all cancer types, *CA9* had the highest average oncogene score on Chromosome 9. *CA9* is a known hypoxia marker and cancer drug target (22), contributes to AKT activation in kidney cancer (23) and is associated with distant metastasis in early-stage cervical cancer if highly expressed (24), supporting its high oncogene score.

Overall our results revealed a highly complex picture of the aberration landscape of human cancer. Integrating different data and cancer types allowed prioritizing a short list of highly relevant candidate cancer genes. We found evidence that frequent focal events highlight probable cancer gene candidates, thus suggesting that the genomic context of aberration patterns provides valuable information.

### Known cancer genes receive high scores

To validate cancer gene prioritization by OncoScape, we analyzed the scores of genes in three types of datasets: 1) frequently mutated cancer genes (4–6), 2) databases of known cancer genes (3, 25) and 3) all human genes and their cancer associated publications. First, we analyzed the prioritization scores of cancer gene candidates frequently mutated in breast or colon cancer (4), across twelve (5) or twenty one (6) different cancer types. The somatic mutation datasets used in our study were also included in the two latter studies. In all cases, the prioritization scores of these genes were significantly higher than the scores of all other genes using the Wilcoxon test (Tables S3-S7). Although these lists against which we validated had been derived using somatic mutation data only, the other data types included in our analysis also showed significantly more aberrations for these genes (Tables S3-S7). OncoScape predicted a number of new candidate oncogene-and tumor suppressor-like genes scoring higher than all but one gene in the previously published lists. For example, we identified the ribosome biogenesis protein BOP1 and RNA-binding protein 39 (RBM39) as new high scoring oncogene candidates. BOP1 has been shown to contribute to tumorigenesis in colorectal cancer (26) and RBM39 has been shown to interact with estrogen receptors and c-JUN (27), which are known for their relevance to cancer.

Second, we investigated the scores of the 513 genes in the Cancer Gene Census (CGC) (2) and found that their aberration scores, calculated as the sum of the oncogene and tumor suppressor scores, were significantly higher than the scores of human genes not included in CGC (Wilcoxon p-value < 2.2*10^−16^, Figure 3a). We also found many additional genes with scores comparable to CGC genes suggesting that these genes are important candidate cancer genes. In total, eleven genes had more extreme combined scores than any CGC gene. These new candidate genes included Bcl-2-like protein 1 (BCL2L1), which is a potent inhibitor of cell death (28). In addition, the 716 genes in the Tumor Suppressor Gene Database (TSGene) (25) had a significantly higher tumor suppressor gene score than other genes (Wilcoxon p-value = 1.95*10^−8^, Figure 3b). These results show that our scoring successfully recovered known cancer genes. Moreover, known tumor suppressor genes were mostly classified as tumor suppressor genes supporting the distinction between activating and inactivating aberrations.

**Figure 3:**
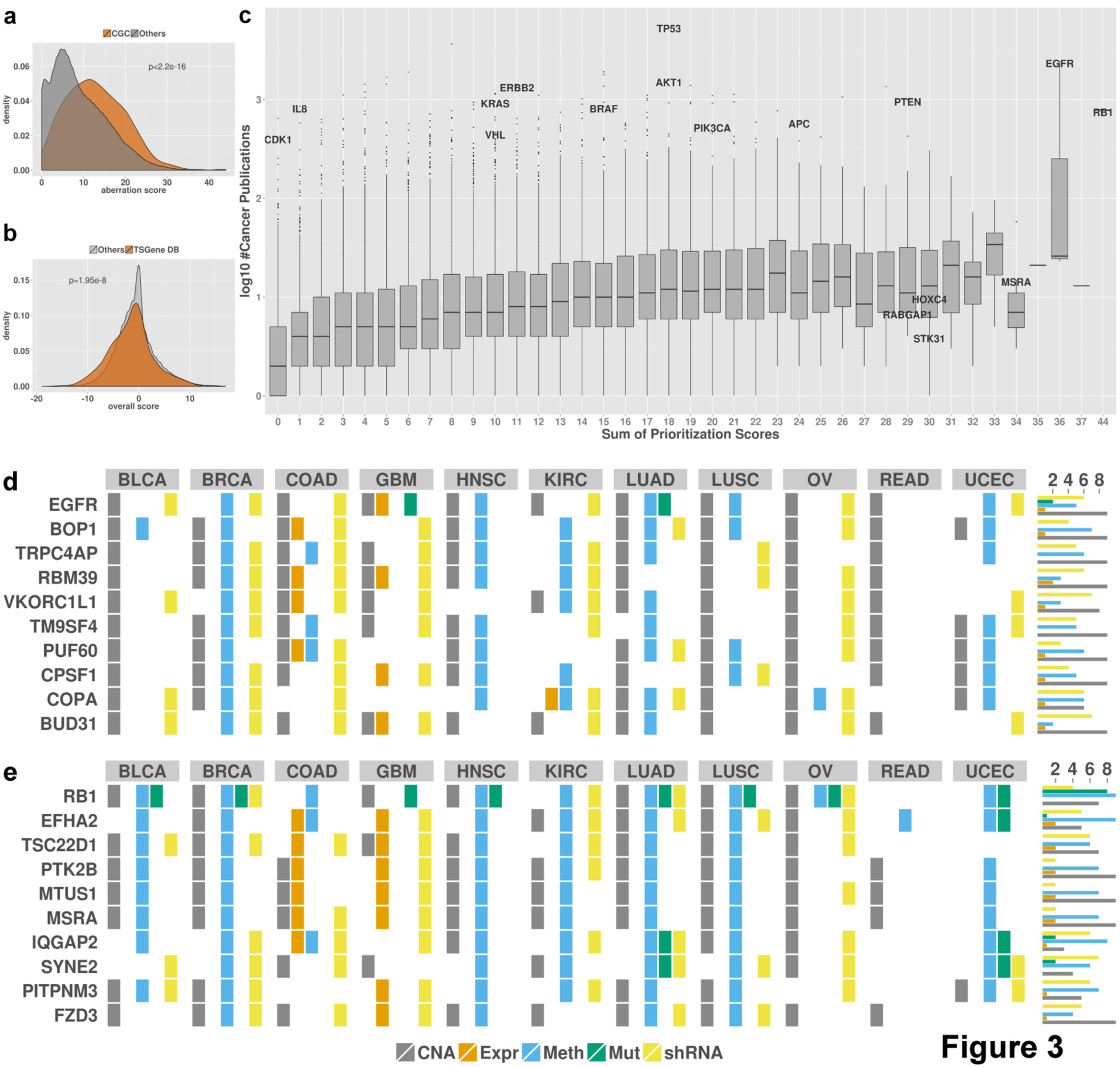
Comparison of prioritization scores with cancer gene databases and publications, and detailed aberration profiles of the highest scoring genes. (a) All human genes are sorted according to the sum of their oncogene and tumor suppressor scores across all cancer types. CGC genes, marked with gray lines, show a trend towards high prioritization scores. (b) All human genes are sorted according to their tumor suppressor scores summed over all cancer types. Known tumor suppressor genes (according to TSGene) were marked and show a significant trend towards prioritization as tumor suppressors. (c) Boxplot comparing prioritization scores with number of cancer-related publications. On the x-axis, genes are categorized using the sum of their oncogene and tumor suppressor gene scores across all cancer types. The y-axis depicts log_10_ of the number of cancer-related publications. Each box summarizes the genes with the same prioritization score. Some well-known cancer genes are labeled. Aberration profiles of the top scoring oncogenes (d) and tumor suppressor genes (e) across all cancer types. Colored boxes indicate that the given gene was aberrated for the respective data type in the associated cancer. Marginal histograms indicate the number of different cancer types with aberrations in the given gene.

Finally, for a global analysis, we utilized the number of cancer-related publications as indicator for a gene’s relevance for cancer development and progression. Overall, we found a good correlation between the aberration score and the number of cancer-related publications (Spearman correlation = 0.37, Figure 3c). This analysis highlighted well-known tumor suppressors and oncogenes with high prioritization scores and a large number of cancer-related publications. These include *RB1* and *EGFR*, the top-scoring tumor suppressor and oncogene, respectively (Figures 3d,e). Furthermore, two particularly interesting groups of genes emerged from this analysis. First, many genes were associated with a large number of cancer-associated publications, but were only rarely aberrated in tumor samples. Such genes included *CDK1*, important for cell cycle regulation, and *IL8*, an important immune system signaling molecule. Second, many genes with high prioritization scores were not frequently described before in the cancer literature. The predicted oncogene *STK31* (Figures 3d,S6), in the top 20 oncogene candidates with a total oncogene score of 20 across the 11 cancer types, is a largely uncharacterized protein kinase. It was mentioned in only five cancer-related publications implicating it as frequently mutated in melanoma (29) and as potential drug target for colorectal cancer (30). *MSRA* scored highly as tumor suppressor gene in several cancer types (Figures 3e,S6), which was supported by its protective role against protein oxidation and the finding that its down-regulation resulted in a more aggressive phenotype in several human cancers (31, 32).

Taken together, these results show that OncoScape successfully recovered known cancer genes while assigning high scores to new genes that likely play an important role in cancer development and progression. We identified a group of genes which had not been frequently mentioned in the cancer-related literature but proved to be highly aberrated across several cancer types. Importantly, several of these genes were not commonly mutated but often targeted by different kinds of aberrations attesting to the advantage of integrating different data types and the power of our approach to identify interesting leads for further studies.

### Context-dependent oncogenes and tumor suppressor genes

The scoring of known tumor suppressor genes demonstrated that our approach allows to distinguish between genes showing oncogene-and tumor suppressor-like behavior. However, we identified potentially activating aberrations for several known tumor suppressor genes. Some genes in TSGene, for instance, had higher oncogene scores than tumor suppressor scores (Figure 3b), which could be explained by tissue type-dependent functions and dual roles for certain genes. *NOTCH1* is one of the genes known to act as oncogene or tumor suppressor in a tissue type-dependent manner (33). Indeed, *NOTCH1* was classified as a tumor suppressor in six tissues (breast invasive carcinoma [BRCA], head and neck squamous cell carcinoma [HNSC], kidney renal clear cell carcinoma [KIRC], lung squamous cell carcinoma [LUSC], ovarian serous cystadenocarcinoma [OV] and uterine corpus endometrial carcinoma [UCEC]) in our analysis and as oncogene in two other tissues (colon adenocarcinoma [COAD] and lung adenocarcinoma [LUAD]). Overall, we found 739 genes indicated as oncogene and tumor suppressor based on at least two different data types for different cancer types. A particularly interesting example was the well-known tumor suppressor gene *RB1*, which was scored as oncogene-like in COAD (Figure S7). This unexpected finding is concordant with previous reports (34, 35) and suggestions that it acts as oncoprotein in colorectal cancer (36). *PUF60*, which ranked among the top ten oncogene candidates across all cancer types (sum of TS scores=7, sum of OG scores=19) (Figure 3d), but in BRCA, its TS score is as high as its OG score. *PUF60* was described to repress *MYC* expression (37), while one specific isoform has been shown to inhibit apoptosis and enable higher expression of *MYC* (38). *PTK2B* was ranked among the ten genes with highest tumor suppressor score (sum of TS scores=20, sum of OG scores=9), mostly because of DNA copy number and methylation changes (Figure 3e), but has also been linked to increased proliferation and invasiveness of hepatocellular carcinoma (39) and MAP kinase pathway activation (40). In KIRC, it received a TS score of 3 and an OG score of 2 indicating conflicting evidence for this cancer type. These examples show that many genes might act as oncogenes or tumor suppressors depending on the cellular context. Our analysis provides a rich resource to identify such cases and the types of aberrations for these genes found in various cancer types. This resource is made available through a web interface (http://oncoscape.nki.nl) where users can explore the best candidates in each cancer type but also annotate a list of genes of interest with tumor suppressor and oncogene results.

### Pathway aberration patterns

Cancer is a disease of pathways, which can be affected by aberrations in different genes (15, 16), for example mutations in *BRAF* and *KRAS* leading to activation of the MAP kinase pathway (41). To facilitate the identification of broader biological categories targeted by aberrations, we calculated pathway scores by averaging combined scores of all genes belonging to one pathway (Figure 4a). Pathway scores for some cancer types were skewed towards higher tumor suppressor scores, for example BRCA and COAD, while in other cancers, like GBM and OV, oncogene pathway scores were generally higher. A similar observation could be made for individual pathways. Calcium signaling, recognized for its importance in cancer (42, 43), was targeted by inactivating aberrations in most genes with the exception of growth factor receptors and PLC-beta isozymes (Figures 4,S8). Aberration profiles of other pathways, such as NF-Kappa B signaling, were clearly dependent on cancer type (Figure S9). Importantly, in some pathways such as PI3K-AKT signaling, individual branches seemed to be targeted consistently across cancer types, while other parts were found to be more heterogeneous (Figure S11).

**Figure 4:**
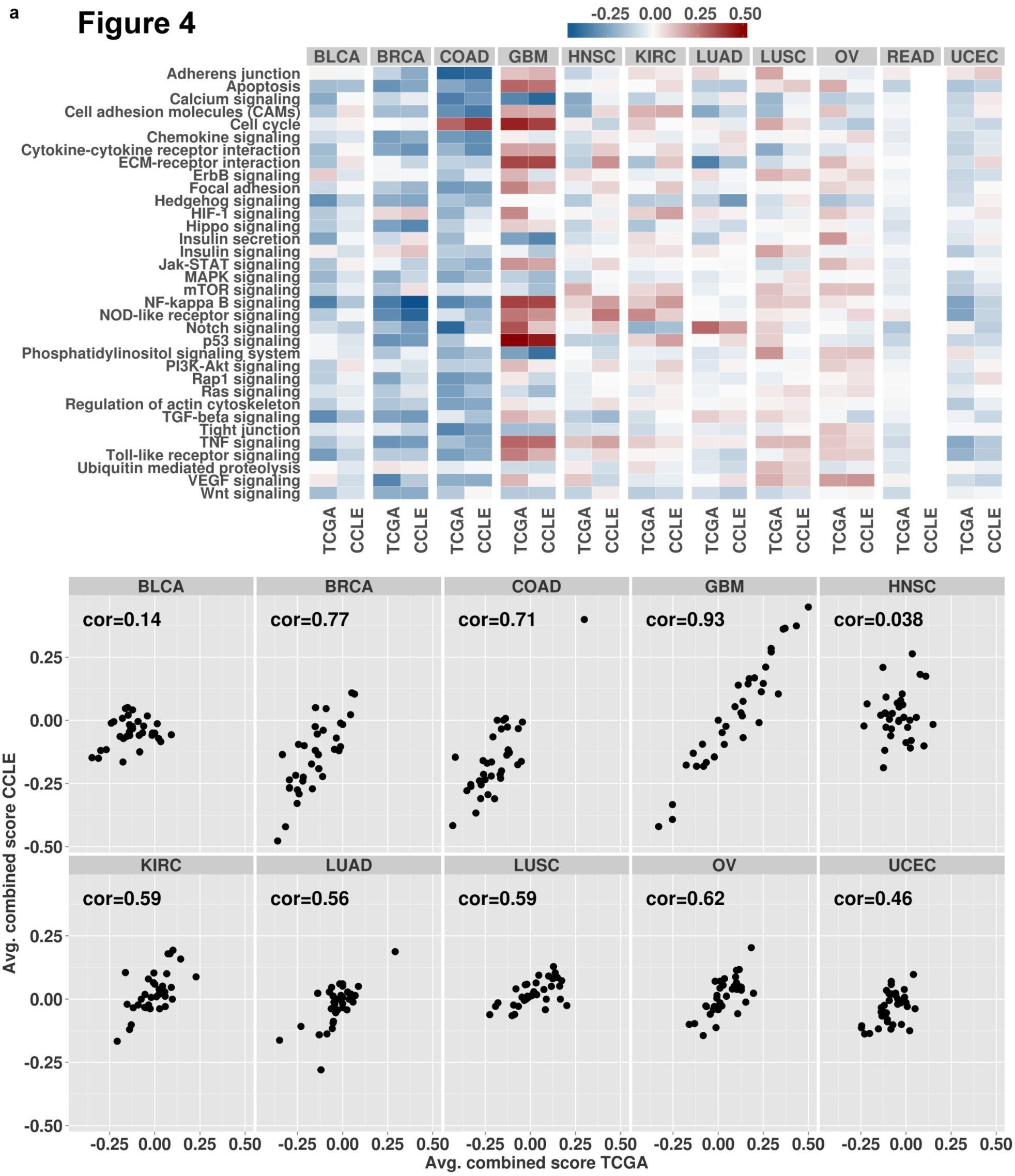
Prioritization scores for 34 selected pathways in tumor samples and cell lines as (a) heatmap and (b) scatter plot. Pathway scores were calculated by averaging scores of all genes belonging to a certain pathway. No rectal cancer cell lines were available. In the scatter plot, each dot represents one pathway. Pathway scores in tumor samples and cell lines are shown on the x-axis and y-axis, respectively. Correlation values are Pearson correlation coefficients between tumor sample and cell line scores.

### Subtypes specific aberrations in colorectal and breast cancer

An interesting application of OncoScape is the identification of genes that are aberrated in specific subtypes of a disease. As an example, we performed an analysis of the recently published four consensus subtypes of colorectal cancer (17) and were able to recapitulate many described characteristics of these subtypes. Guinney and colleagues described consensus subtypes CMS1 and CMS3 as having few copy number changes while samples in CMS2 had many copy number alterations. In line with this, OncoScape scored no genes for copy number changes in CMS1 and only 34 genes in CMS3. In contrast, copy number changes were identified for more than 2100 genes in CMS2, including amplifications of HNF4A as described previously. We also identified the mutations in BRAF as being specific for CMS1. CMS4 was described as having altered TGF-beta signaling, which was the pathway with the highest OncoScape pathway score for this subtype. Importantly, we could also identify new subtype-specific events. OncoScape found the serine/threonine-protein kinase Nek3 to be targeted by loss of DNA methylation and copy number gains unique to CMS2. NEK3 has been described to regulate motility of breast cancer cells and to be overexpressed in breast cancer tissue (44). Our results suggest it as being deregulated in a specific suptype of colorectal cancer, which could represent a specific vulnerability for a subset of tumors. These results show that OncoScape can identify aberrations that are specific to individual subtypes and that could be targeted in this patient subpopulation.

OncoScape was also used to identify drivers of two subtypes of invasive lobular breast cancer: the Hormone Related (HR) subtype and the Immune Related (IR) subtype (45). Michaut and colleagues found four potential drivers of the HR subtype: PGR, GATA3 and FN1, were up-regulated at the mRNA and RPPA level. Similarly, YAP1 was deleted and down-regulated in HR, suggesting a possible role as driver of this subtype.

### Comparing cell lines to primary tumors

As cancer cell lines are important models for studying the biology of cancer, we compared the molecular aberration landscape of cancer cell lines to that of tumor samples. To this end, we mapped cancer cell lines from the Cancer Cell Line Encyclopedia (CCLE) (10) based on their tissue of origin to the eleven TCGA cancer types included in our analysis. We performed the analysis using the 327 cancer cell lines that could be mapped to one of these cancer types (see Methods for details). Several differences between the datasets have to be considered when interpreting the results of this comparison. Rectal cancer was not represented in the cell line set, leaving 10 cancer types for the comparison. Since no matched normal samples were available for cell lines, we compared cell lines to tissue-matched normal samples from TCGA. DNA methylation data were not available for the cell lines and the mutation data were limited to about 1,650 genes.

The overall number of aberrated genes was similar between cell lines and tumor samples, as was the number of genes with aberrations in the different data types (Figure S11). Few genes were found frequently mutated in the cell lines and a substantially higher number of genes were affected by DNA copy number alterations and gene expression changes, similar to the observations in tumors. Significantly altered gene expression was especially common in GBM and COAD, as observed before for tumor samples. With the exception of HNSC, genes with maximal scores in tumors were also found to be aberrated in cancer cell lines (Table 2), although the obtained scores were generally lower in cell lines. These results suggest that individual genes generally show comparable aberration profiles in cell line panels of most cancer types analyzed here. The comparison of pathway scores between cell lines and tumors revealed large differences between cancer types (Figure 4). We found a high correlation between pathway scores for instance for GBM, BRCA and COAD indicating that the available cell lines represented the set of tumors very well. Other cell line panels showed little correspondence with tumor samples, most notably HNSC and BLCA.

Our results stress the importance of carefully selecting an *in vitro* model system. Cell lines of several cancer types, such as GBM, BRCA and COAD, have a high overall correspondence with tumor samples implicating a good coverage of the tumor aberration space. Importantly, our analysis highlighted several cancer types (HNSC and BLCA, in particular) and pathways with only little similarity between tumors and cell lines which demands greater caution when choosing particular cell lines as model systems.

### OncoScape web portal

We implemented a web portal (http://oncoscape.nki.nl) for exploring OncoScape results interactively. It allows accessing detailed scoring results for all cancer types and visualizing them in different types of plots. A pre-defined gene list can also be uploaded and annotated with oncogene and tumor suppressor gene prediction scores in all cancer types. Additionally, the web portal provides functionality for directly comparing results from tumor samples and cell lines. All source code can be downloaded from GitHub (https://github.com/andreas-schlicker/OncoScape); result files can be accessed through Sage Bionetwork’s Synapse (doi:10.7303/syn2518467).

## DISCUSSION

We presented OncoScape, a new method for integrating several genome-wide datasets in order to characterize the molecular aberration landscape of human cancer. Many identified genes were rarely mutated but frequently affected by other aberrations such as DNA copy number alterations or methylation changes. These genes would not have been detected by methods that rely solely on somatic mutations, thus proving the value of integrating different genome-wide datasets. In contrast to most previous approaches, we prioritized genes as potential oncogenes or tumor suppressor genes based on the type of aberrations found. It should be borne in mind that we prioritize genes based on a score aggregated across different data types. This scoring system favors genes that show aberrations in multiple data types over genes that are almost exclusively aberrated in a single data type. Therefore, we would not identify genes that are often mutated but do not show any other alterations. The analysis of aberration scores across pathways highlighted some pathways, like calcium signaling, activated or inactivated across all cancer types, which could implicate them as targets for therapeutic intervention in many different cancer types. Others exhibited tissue-specific aberration patterns suggesting that corresponding compounds would have tissue-specific activity.

By analyzing data for available cancer cell lines of 10 cancer types, we provide the first large-scale comparison of the molecular aberration landscapes of tumors and cell lines. Although data availability slightly differed for tumors and cell lines, the overall characteristics of the results indicate that they were comparable. We did not perform a comparison for single cell lines as presented previously for ovarian cancer (14). Overall, we found a good similarity between the aberration landscapes of ovarian cell lines and tumors, but we also observed several pathways with apparent differences, including TGF-beta signaling and insulin secretion. Aberrations found in the panel of colon cancer cell lines closely resembled colon tumors, which is in line with previous results (13). Our analysis also highlighted some cancer types for which available cell lines and tumor samples differ to a large degree, notably HNSC and BLCA. For bladder cancer, our results agree with recent findings published by Earl *et al*. [10.1186/s12864-015-1450-3]. They identified significant differences in the mutation rates of important oncogenes and tumor suppressor genes between tumors and cell lines. In their analysis of gene expression subtypes, Earl and colleagues noted that while aggressive subtypes are not that common in patients they were predominant in the analyzed bladder cancer cell lines. Together with our results, this stresses the importance of carefully selecting cell lines as *in vitro* models for investigation of these cancer types.

In summary, we provide a comprehensive overview of the molecular aberration landscape of tumors of 11 cancer types and cell lines of 10 cancer types obtained using our new method OncoScape. Our results stress the value of integrating different data types when identifying candidate cancer genes and suggest many new leads for further investigation. The comparison between tumors and cell lines provides a valuable resource for researchers to guide the selection of cell lines for research into mechanisms of oncogenesis as well as for the development of new therapeutics. Importantly, OncoScape can be applied in many additional contexts, such as the comparison of patients responding or being resistant to specific treatments or to identify specific characteristics of different subtypes of a certain disease.

---

Table 1: Number of available primary tumor and matched normal samples per cancer type. Each row contains the number of samples available for the specified data type. Rows 5 and 6 contain the number of samples with available mRNA expression data and copy-number data or DNA methylation data, respectively. The last row contains the number of cell lines available in CCLE for each cancer type.

Table 2: Cell line aberration status of the highest scoring genes for tumor samples. Genes are classified as aberrated in cell lines if they received an oncogene or tumor suppressor score of at least one.

